# Executive Dysfunctions In The EAAT3 Overexpressing Mouse Model Of Compulsive Behavior

**DOI:** 10.64898/2026.09.03.749220

**Authors:** F. Henríquez-Belmar, M. Churruca-Muñoz, W. Plaza-Briceño, P. Acevedo-Hernández, M.J. Cossio, A.E. Chávez, J.P. Casanova, P.R. Moya

## Abstract

Executive functions are a set of cognitive processes that regulate behavior and adapt responses to changing demands. Impairments in these processes are common in various neuropsychiatric conditions, including obsessive-compulsive disorder. Mice overexpressing the neuronal glutamate transporter EAAT3 in the forebrain (EAAT3glo/CaMKII) exhibit increased compulsive behaviors and synaptic alterations relevant to OCD. However, whether EAAT3 overexpression affects executive functioning across multiple cognitive domains has not been studied. Using operant conditioning and visuospatial learning tasks, we evaluated the performance of EAAT3glo/CaMKII mice across three behavioral domains commonly associated with executive function: cognitive flexibility, working memory, and inhibitory control. EAAT3glo/CaMKII mice showed impaired cognitive flexibility in both operant extinction and reversal learning. EAAT3glo/CaMKII mice also failed to acquire the Trial-unique, delayed nonmatching-to-location task, suggesting impaired working memory, and showed increased impulsivity in the five-choice serial reaction time task, as evidenced by elevated premature responses. Perseverative responses, response accuracy, and attentional performance were unaffected in EAAT3glo/CaMKII mice. Collectively, these findings suggest that EAAT3 overexpression in forebrain neurons is associated with a cognitive profile characterized by difficulties in flexibility and task acquisition, alongside increased impulsivity, while attentional performance and motivational measures remain intact in a model of compulsive behavior.

## INTRODUCTION

Obsessive-compulsive disorder (OCD) is a neuropsychiatric disorder characterized by intrusive, unwanted thoughts (obsessions), repetitive ritualistic behaviors (compulsions) and anxiety (1). Cognitive deficits are frequently reported in individuals with OCD, and neuroimaging evidence indicates abnormalities in prefrontal-striatal circuits that support executive functions (2). Executive functions are a set of general-purpose cognitive abilities that, by regulating lower-level cognitive processes (e.g., perception, motor responses), allow individuals to regulate their thoughts and behavior and to achieve internally represented goals (3, 4). To date, three primary executive functions have been described: inhibitory control, working memory and cognitive flexibility; these functions operate in an interrelated manner (4). Higher-order executive functions can be derived from these core operations, including reasoning, problem-solving, and planning (5, 6). Consequently, alterations in executive functions can seriously impact daily functioning, and may be important contributors to neuropsychiatric disorders including OCD, where impairments in all cores executive functions have been documented (7–12).

Animal models are valuable tools to study the neurobiological mechanisms underlying psychiatric conditions, including OCD (13–15). Previously, we reported that mice overexpressing the neuronal glutamate transporter EAAT3 (EAAT3glo/CaMKII) exhibit increased compulsive and anxiety-like behaviors, as well as impaired long-term extinction of fear conditioning (16). EAAT3glo/CaMKII mice have also alterations in glutamatergic cortico-striatal transmission and NMDA receptor subunit composition (16). EAAT3 is an important regulator of glutamatergic transmission by controlling glutamate spillover at synaptic level (17). Glutamate is a neurotransmitter critical in brain regions controlling executive functions (4, 18) and its dysregulation may contribute to the pathophysiology of OCD (19, 20). However, the impact of EAAT3 overexpression in executive functions has not yet been thoroughly studied in EAAT3glo/CaMKII mice. Therefore, we evaluated the performance of EAAT3glo/CaMKII mice in cognitive flexibility, working memory, and inhibitory control using operant conditioning and visuospatial learning tasks. Understanding whether EAAT3 overexpression selectively or broadly affects executive function could provide insight into the specific cognitive domains vulnerable to glutamatergic dysregulation and help define the translational relevance of this model to human compulsive disorders.

## MATERIALS AND METHODS

### Animals

All experimental procedures were approved by the Institutional Bioethics Committee of the Universidad de Valparaíso (BEA 131-19). 10-week-old male and female control (EAAT3glo) and EAAT3glo/CaMKII mice were used (16). Mice were weaned on postnatal day 21 and housed in groups of four to six animals per cage, with controlled temperature (21 ± 2°C) and humidity (40-70%), in a 12-hour light/dark cycle (lights on at 8:00 AM) and had access to water and food *ad libitum*. All experiments were performed blind to mouse genotype.

### Behavioral tests

#### Mild food restriction protocol

We followed the protocol outlined by Horner et al. (21). Mice were housed in pairs per cage and provided with 2-3 g of food per animal daily. Their weight was recorded daily to ensure that they lost less than 1 g per day, adjusting the amount of food provided, and *ad libitum* access to water. Operant conditioning tests were initiated when mice reached 85% of their initial weight. The animals were kept on food restriction throughout the tests, and their weight was monitored weekly.

#### Standard operant conditioning

We followed a standard nose-poke operant conditioning protocol carried out in an open-source operant chamber (22) with three holes (left, center, and right) equipped with photo-interrupter sensors controlled by an Arduino microcomputer. Briefly, mice are required to learn that the active hole is associated with the delivery of a reward (10 μL of 30% sucrose) in the center hole through a calibrated syringe. Before the experiments began, mice were habituated to the reward by placing a small dish containing 3 mL of 30% sucrose solution in the home cage for two consecutive days. Mice were then habituated to the behavioral room and the operant chamber, without any training. After habituation, mice were placed in the operant boxes for one hour daily for nine consecutive days (Training phase), during which the responses made in the active, inactive, and central holes were recorded. Mice were trained one at a time. Behavior was considered acquired if the following criteria were met: *1)* obtaining more than 20 sucrose deliveries in three consecutive sessions and *2)* exhibiting a discrimination ratio greater than 3:1 in active versus inactive responses for three consecutive days (22). Once the training protocol had been completed, and only if the acquisition criteria had been met, mice were separated into two groups to undergo either ‘extinction’ or ‘reversal’ protocol during the Test phase. To evaluate extinction, mice were placed in the operant box for one hour daily for six days, with the active hole no longer providing a reward. For reversal learning, the reward-hole associations were reversed, meaning that the animals had to extinguish the previously rewarded response and learn to choose the previously unrewarded, now rewarded option. This protocol was performed for four consecutive days (22).

#### Evaluation of Inhibitory control using the Five-choice serial reaction time task (5-CSRTT)

The 5-CSRTT (23) was performed in Bussey-Saksida operant chambers (Campden Instruments, Loughborough, LE, UK) to evaluate visuospatial attention, compulsivity and motor impulsivity according to the protocol outlined by Mar et al. (24). The 5-CSRTT comprises three consecutive phases:

a) Preparation phase: Once mice reached 85% of their original weight (see above), they were habituated to the liquid reward (chocolate milk) for 3 days in their home cage.

b) Pre-training phase: This phase consists of six stages, four habituations lasting 1-2 days, plus two stages introducing the mice to use the screens (Supplementary Table 1). During this phase, mice gradually become used to the operant chambers, the reward, and the stimuli while progressively learning the behavior of touching the screen appropriately when a stimulus is presented.

c) Testing phase: This phase consists of 11 stages, each lasting 7-10 days, in which the animal is expected to maintain and divide its attention among the five locations lined up on the screen to detect and respond to a visual stimulus that progressively becomes shorter, as described in Supplementary Table 1. The flowchart for this stage is shown in Figure S1. Once the animal achieves a stable baseline in the last step of this phase for two consecutive days, its performance in the basic 5-CSRTT can be analyzed. Several aspects of executive function were evaluated:

- Response Accuracy (% accuracy): Refers to the number of correct responses divided by the total number of both correct and incorrect responses. This value is the primary index of divided and sustained attention.

- Omissions (% omission): Refers to the number of trials in which the mouse did not respond divided by the total number of completed trials. The percentage of omissions reflects overall attention.

- Premature Responses: Refers to the number of responses made during the intertrial interval (ITI). This is the primary measure of impulsive behavior.

- Perseverative Responses: Refers to the number of responses (nose pokes) made in any opening after the mouse gives a correct response but before retrieving the reward. This is a measure of compulsive behavior.

- Magazine Entries: Refers to the number of nose-poke responses in the reward receptacle. This is a measure of the mouse’s motivation.

- Reward Retrieval Latency: This is the average time it takes the mouse to retrieve the reward. This parameter is related to the mouse’s motivation to perform the task.

#### Evaluation of working memory using the Trial Unique Non-Matching to Location (TUNL) Task

The TUNL is a working memory task based on the delayed matching or non-matching-to-position paradigm. Briefly, the predefined sample and the novel choice location can be randomly selected from multiple response locations and vary between trials within a session, making this task “trial-unique”. Specifically, the requirement to retain spatial information during the delay can be reduced by taking advantage of the predictability of the correct location. The TUNL task avoids this issue by using a variety of spatial locations, making the correct choice location less predictable (25). The TUNL protocol consists of four main phases:

a) Pre-training phase: Including food restriction, weight monitoring, and reward habituation stages as described above for the 5-CSRTT.

b) Training phase: Includes the same 6 stages described in Supplementary Table 1 for the 5-CSRTT and a final stage, in which the animal is presented with the task and punished when performing an incorrect trial. The punishment consists of a reinforcement trial that is repeated until the animal completes the test correctly. More specifically, this task consists of two phases. In the base phase, a stimulus (white square) is presented to the animal at one of the possible locations on the screen; once the mouse touches the stimulus, it disappears from the screen. After a variable delay period, the test phase begins in which the old stimulus or test stimulus is presented along with a new stimulus in a new location. The animal must touch the new stimulus to make a correct response.

c) Acquisition: Consists of 2 stages (described in Supplementary Table 2), each of which is composed of 3 sub-stages lasting between 10-14 days each. Briefly, Stage 1 requires that sample and test stimuli be presented in any location except the center (square 3) and with different levels of spatial separation or gap (G1, G2, and G3), while Stage 2 requires that one of the two stimuli be presented in the center with different levels of spatial separation or gap (G1, G0, and G1&0).

#### Evaluation of working memory using the spontaneous alternation test in a T-Maze

The spontaneous alternation test in a T-maze is based on rodentś natural tendency to prefer exploring novel areas over familiar ones, leading them to alternate between target arms (26). Mice were acclimated for one hour in the behavior room and then carefully placed at the starting arm, facing the back wall. Once an animal fully entered one of the two arms (tail criterion), it was confined to the chosen arm for 30 seconds using a door. This procedure was repeated for six additional trials, recording the time it took the animal to choose and enter the arm. If the chosen arm differs from the previous one, it is considered an alternation.

### Statistical Analysis

Statistical analyses were performed using GraphPad Prism 8.4.3 (GraphPad Software, San Diego, CA, USA). Data normality was assessed using the Shapiro-Wilk test. Outliers were identified and removed using Grubbs’ sequential test (α = 0.05). For acquisition, extinction, and reversal learning data, between-genotype differences in behavioral trajectories were assessed using mixed-effects repeated-measures models (REML), with Session as the within-subject factor and Genotype as the between-subjects factor. Active hole, inactive hole, and magazine responses were analyzed separately. *Post-hoc* comparisons between genotypes at individual sessions were corrected using Sidak’s method. Area under the curve (AUC) was calculated using the trapezoid rule and compared between genotypes using unpaired t-tests or Mann-Whitney U tests.

## RESULTS

Both male and female mice were included in all experiments. Based on previous characterization of this mouse model showing no sex differences in compulsive-like phenotypes (16), data from both sexes were pooled for all analyses reported here.

### EAAT3glo/CaMKII mice show normal acquisition but impaired extinction of operant learning

Control EAAT3glo (n=22) and EAAT3glo/CaMKII (n=25) mice were subjected to a simple operant learning task. Both groups correctly acquired operant behavior, reaching a 3:1 ratio of active to inactive nose pokes by the second day of training (Fig. 1A). Active hole responses progressively increased across sessions while inactive hole responses decreased, with no between-genotype differences in learning trajectories (Mixed-effects REML, Session × Genotype: active F(8,182)=0.671, p=0.715; inactive F(8,397)=0.654, p=0.732). At the end of the protocol, both groups showed significantly more active hole responses at day 9 compared to day 1 (Fig. 1B, p<0.001), and significantly fewer inactive hole responses (Fig. 1C, p<0.0001), with no differences in magazine responses between genotypes (Fig. 1D, Session × Genotype: F(8,200)=0.791, p=0.611). These results indicate that EAAT3 overexpression does not alter the acquisition of operant behavior.

**Figure 1.**
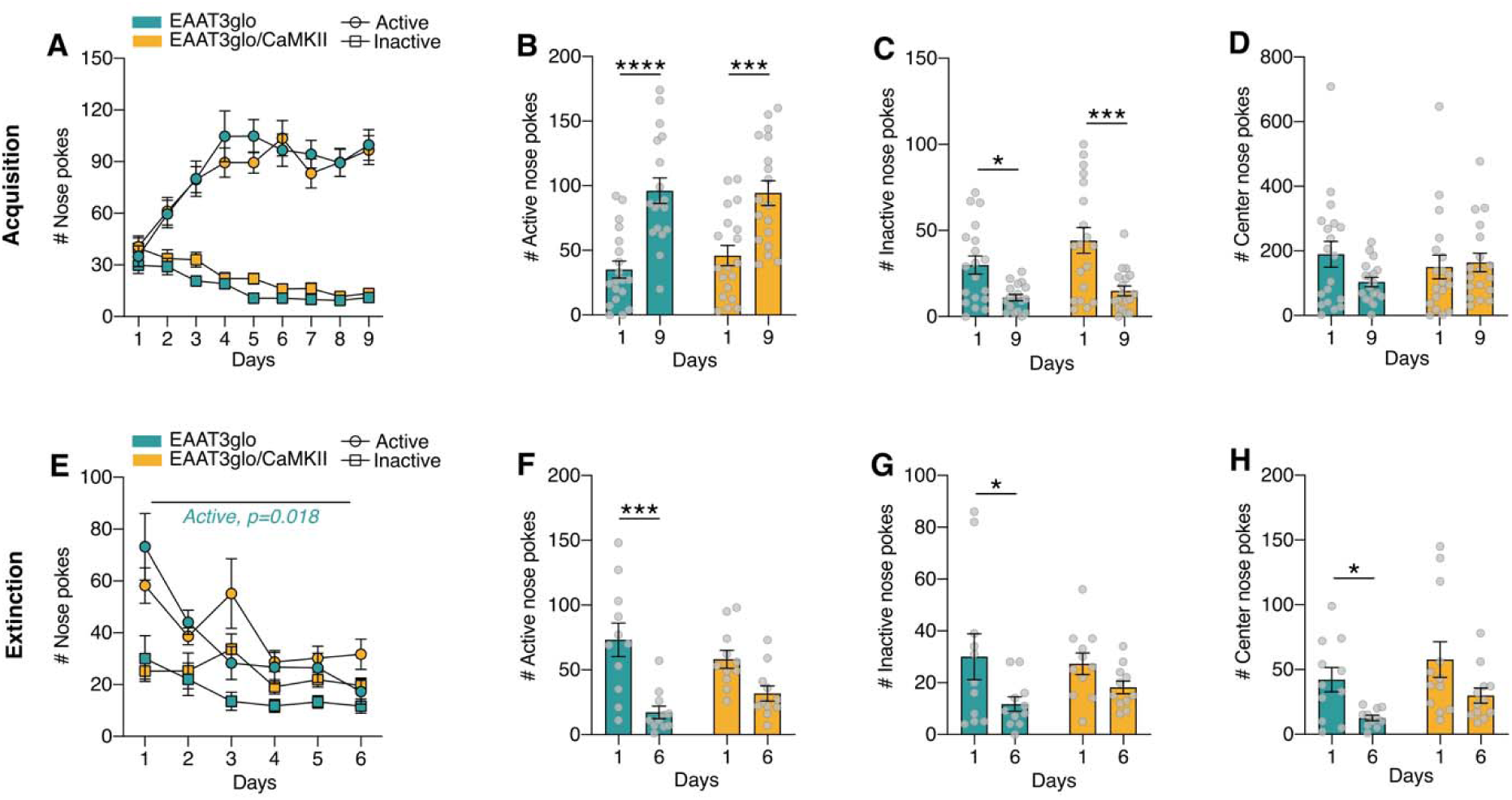
EAAT3glo/CaMKII mice show impaired extinction of operant learning. (A) Active (circles) and inactive (squares) hole responses across nine acquisition sessions. Both genotypes learned the task at the same rate, with no between-genotype differences in response trajectories (Mixed-effects REML, Session × Genotype: active F(8,182)=0.671, p=0.715; inactive F(8,397)=0.654, p=0.732). (B) Active hole responses at day 1 vs day 9. Both genotypes showed a significant increase in active responding across training (EAAT3glo p<0.0001; EAAT3glo/CaMKII p<0.001). (C) Inactive hole responses at day 1 vs day 9. EAAT3glo mice showed a significant decrease (p<0.05); EAAT3glo/CaMKII mice showed a significant decrease at a higher threshold (p<0.001). (D) Center hole responses at day 1 vs day 9, no significant differences. (E) Active and inactive hole responses across six extinction sessions. EAAT3glo mice reduced active responding significantly more than EAAT3glo/CaMKII mice (Session × Genotype: F(5,100)=2.884, p=0.018). (F) Active hole responses at day 1 vs day 6. EAAT3glo mice showed a significant reduction (p<0.001); EAAT3glo/CaMKII mice did not. (G) Inactive hole responses at day 1 vs day 6. EAAT3glo mice showed a significant reduction (p<0.05); EAAT3glo/CaMKII mice did not. (H) Center hole responses at day 1 vs day 6. EAAT3glo mice showed a significant reduction (p<0.05); no significant change was observed in EAAT3glo/CaMKII mice. Between-genotype comparison of center responses across sessions was not significant (Session × Genotype: F(5,105)=0.405, p=0.831), ruling out motivational differences. EAAT3glo n=22, EAAT3glo/CaMKII n=25 (acquisition); EAAT3glo n=11, EAAT3glo/CaMKII n=11 (extinction). Mean ± SEM with individual data points. *p<0.05; ***p<0.001; ****p<0.0001.

Following acquisition, EAAT3glo/CaMKII (n=11) and control EAAT3glo (n=11) mice were subjected to extinction by disabling reward delivery. Mixed-effects REML analysis revealed a significant Session × Genotype interaction for active hole responses (F(5,100)=2.884, p=0.018), indicating that the two genotypes followed different extinction trajectories (Fig. 1E). Control mice progressively reduced active hole responding across sessions (Fig. 1F, day 1 vs day 6, p<0.0001), while EAAT3glo/CaMKII mice did not show a comparable reduction. A similar trend was observed for inactive hole responses (Session × Genotype: F(5,50)=2.106, p=0.080), though this did not reach statistical significance (Fig. 1G). Importantly, magazine responses decreased similarly in both groups (Session × Genotype: F(5,105)=0.405, p=0.831, Fig. 1H), ruling out motivational differences as an explanation for the extinction deficit. These results indicate that EAAT3 overexpression impairs extinction of operant behavior.

### EAAT3glo/CaMKII mice show impaired reversal learning

To evaluate cognitive flexibility, EAAT3glo/CaMKII (n=11) and control EAAT3glo (n=11) mice were subjected to a reversal learning protocol in which the previously rewarded and unrewarded holes were switched. Mixed-effects REML analysis revealed significant Session × Genotype interactions for both the previously rewarded hole (-CS: F(3,83)=3.398, p=0.023, Fig. 2A) and the newly rewarded hole (+CS: F(3,83)=3.509, p=0.020, Fig. 2C), indicating divergent behavioral trajectories between genotypes across the reversal sessions. Control mice progressively reduced responding to the -CS hole and increased responding to the +CS hole, while EAAT3glo/CaMKII mice failed to show this pattern. AUC analysis confirmed a significant between-genotype difference for +CS responses (Mann-Whitney, p=0.038, Fig. 2D), but not for - CS responses (Mann-Whitney, p=0.182, Fig. 2B), consistent with a gradual divergence more readily captured by the longitudinal analysis. *Post-hoc* Sidak comparisons between genotypes at individual sessions did not reach statistical significance, reflecting the gradual nature of the effect. Magazine responses did not differ between genotypes across sessions (Session × Genotype: F(3,66)=0.226, p=0.878, Fig. 2E-F), ruling out motivational differences. These results indicate that EAAT3 overexpression impairs cognitive flexibility.

**Figure 2.**
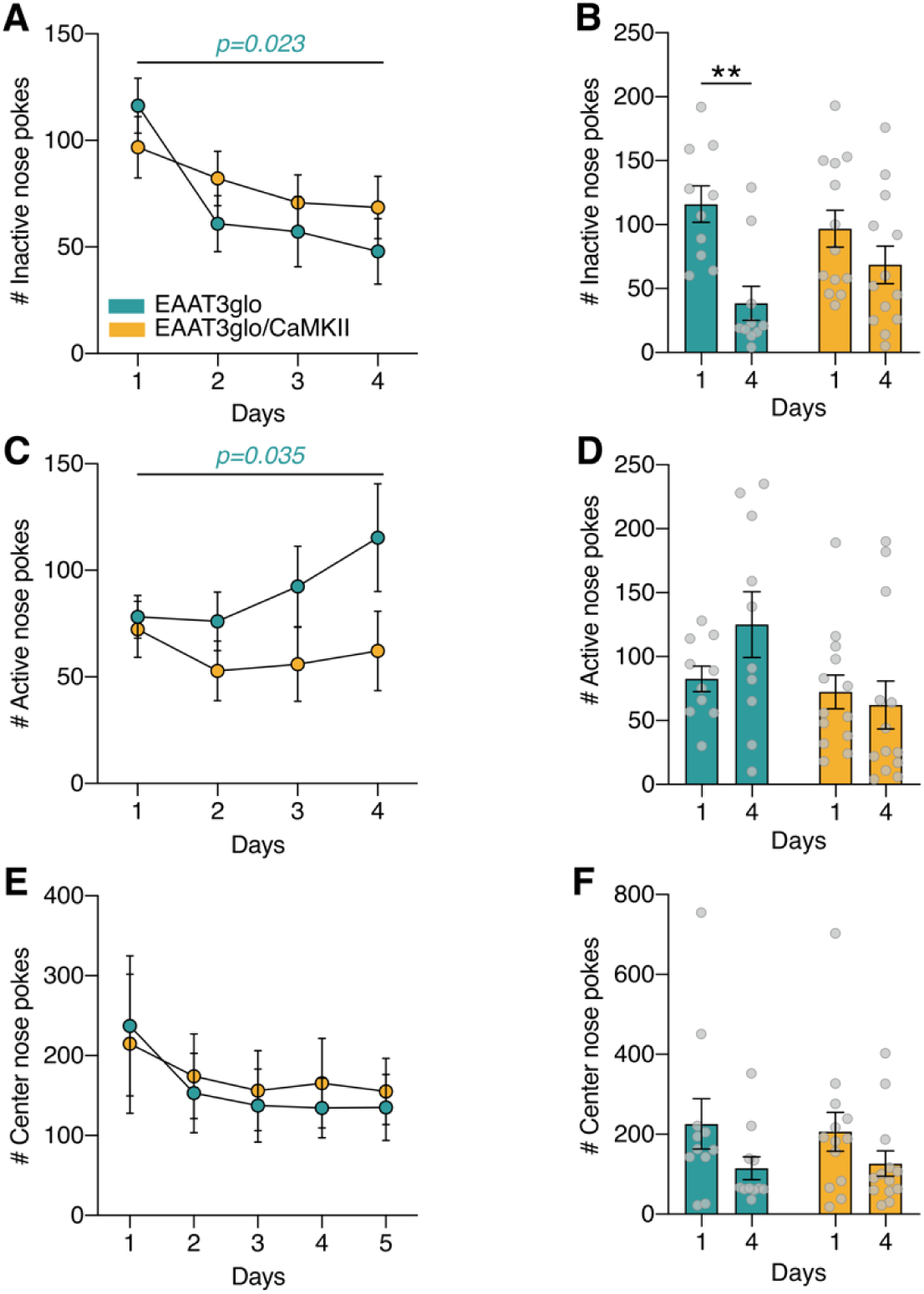
EAAT3glo/CaMKII mice show impaired reversal learning. (A, C) -CS and +CS hole responses across four reversal sessions. EAAT3glo mice reduced responding to the previously rewarded hole (-CS) and increased responding to the newly rewarded hole (+CS). EAAT3glo/CaMKII mice failed to show this pattern (Session × Genotype: - CS F(3,83)=3.398, p=0.023; +CS F(3,83)=3.509, p=0.020). (B, D) Responses at day 1 vs day 4 per genotype. AUC analysis confirmed a significant difference for +CS responses (Mann-Whitney, p=0.038) but not for -CS (p=0.182), consistent with a gradual divergence between trajectories. Post-hoc Sidak comparisons at individual sessions were not significant. (E, F) Magazine responses did not differ between genotypes (Session × Genotype: F(3,66)=0.226, p=0.878; AUC p=0.451), ruling out motivational differences. EAAT3glo n=11, EAAT3glo/CaMKII n=11. Mean ± SEM. *p<0.05.

### EAAT3glo/CaMKII mice show increased impulsivity in the 5-CSRTT

To evaluate inhibitory control and attentional performance, EAAT3glo/CaMKII (n=6) and control EAAT3glo (n=8) mice were tested in the 5-CSRTT (Fig. 3A). No between-genotype differences were found in response accuracy (Mann-Whitney, p=0.77, Fig. 3B), omission rate (Mann-Whitney, p=0.93, Fig. 3C), perseverative responses (Mann-Whitney, p=0.92, Fig. 3D), reward collection latency (p=0.55. Fig. S4A), time-out touches (p=0.48, Fig S4B), or feed-tray panel pushes (p=0.61, Fig. S4C), indicating equivalent motivation, attentional performance, and general motor activity across groups. However, EAAT3glo/CaMKII mice showed a significant increase in premature responses compared to controls (Welch’s t-test: t(6.94)=2.670, p=0.032, Fig. 3E), indicating elevated impulsivity; Welch’s correction was applied due to unequal variances between groups (F-test: p=0.010). Next, to further challenge attentional and inhibitory control processes, mice were subjected to three additional one-day tests with variable stimulus duration and inter-trial intervals (ITI). No between-genotype differences were found in any of these challenge conditions (Figs. S1-S3), suggesting that increased premature responses observed at baseline reflects a specific alteration in impulsivity rather than a broader deficit in attention or inhibitory control.

**Figure 3.**
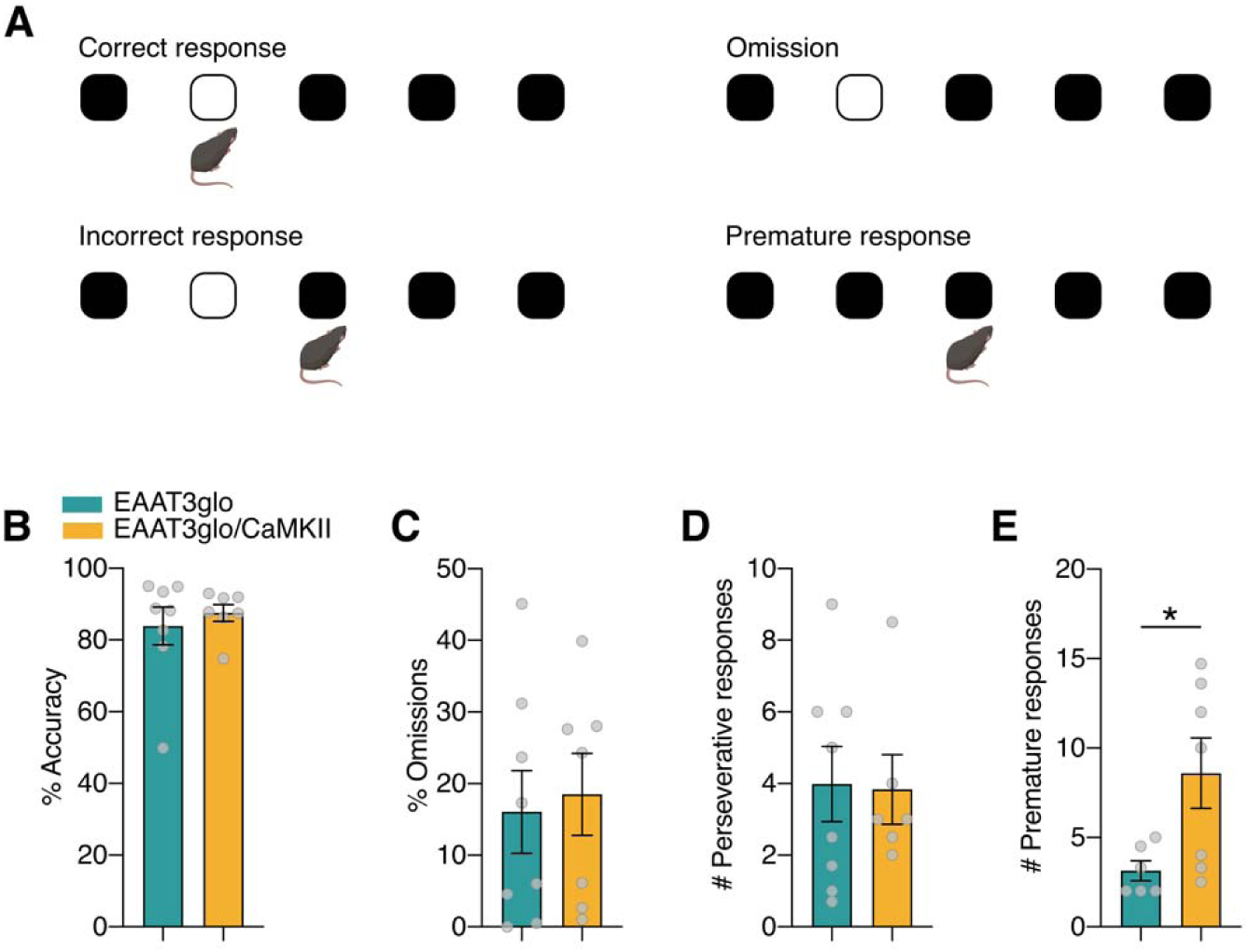
EAAT3glo/CaMKII mice show increased impulsivity in the 5-CSRTT. (A) Task schematic. No differences between genotypes were found in response accuracy (B; p=0.77), omissions (C; p=0.93), perseverative responses (D; p=0.92), reward collection latency (E; p=0.55), time-out touches (F; p=0.48), or feed tray panel pushes (G; p=0.61). EAAT3glo/CaMKII mice made significantly more premature responses (H; Welch’s t-test: t(6.94)=2.670, p=0.032), indicating higher impulsivity. Welch’s correction was used due to unequal variances (F-test: p=0.010). EAAT3glo n=8, EAAT3glo/CaMKII n=6. Mean ± SEM with individual data points. *p<0.05.

### EAAT3glo/CaMKII mice show working memory deficits

We evaluated working memory using two complementary approaches: the TUNL touchscreen task and the T-maze spontaneous alternation test. In the TUNL task (Fig. 4A), EAAT3glo/CaMKII mice (n=7) showed progressive failure to advance through the acquisition stages compared to control EAAT3glo mice (n=7). While all control mice completed every stage of the protocol (100%), EAAT3glo/CaMKII mice showed progressive dropout from training onwards, with none reaching stages S2S0 or S2S0&1. Log-rank (Mantel-Cox) analysis confirmed that the two groups differed significantly in their acquisition trajectories (χ²(1)=15.99, p<0.0001, Fig. 4B). the number of sessions required to reach criterion tended to be higher in EAAT3glo/CaMKII mice at the training stage (Mann-Whitney, p=0.063) and at S1G3 (p=0.391), though these comparisons did not reach statistical significance, likely reflecting the limited sample sizes resulting from progressive dropout (Fig. 4C). These results indicate that EAAT3glo/CaMKII mice are unable to acquire the cognitive demands of the TUNL task.

**Figure 4.**
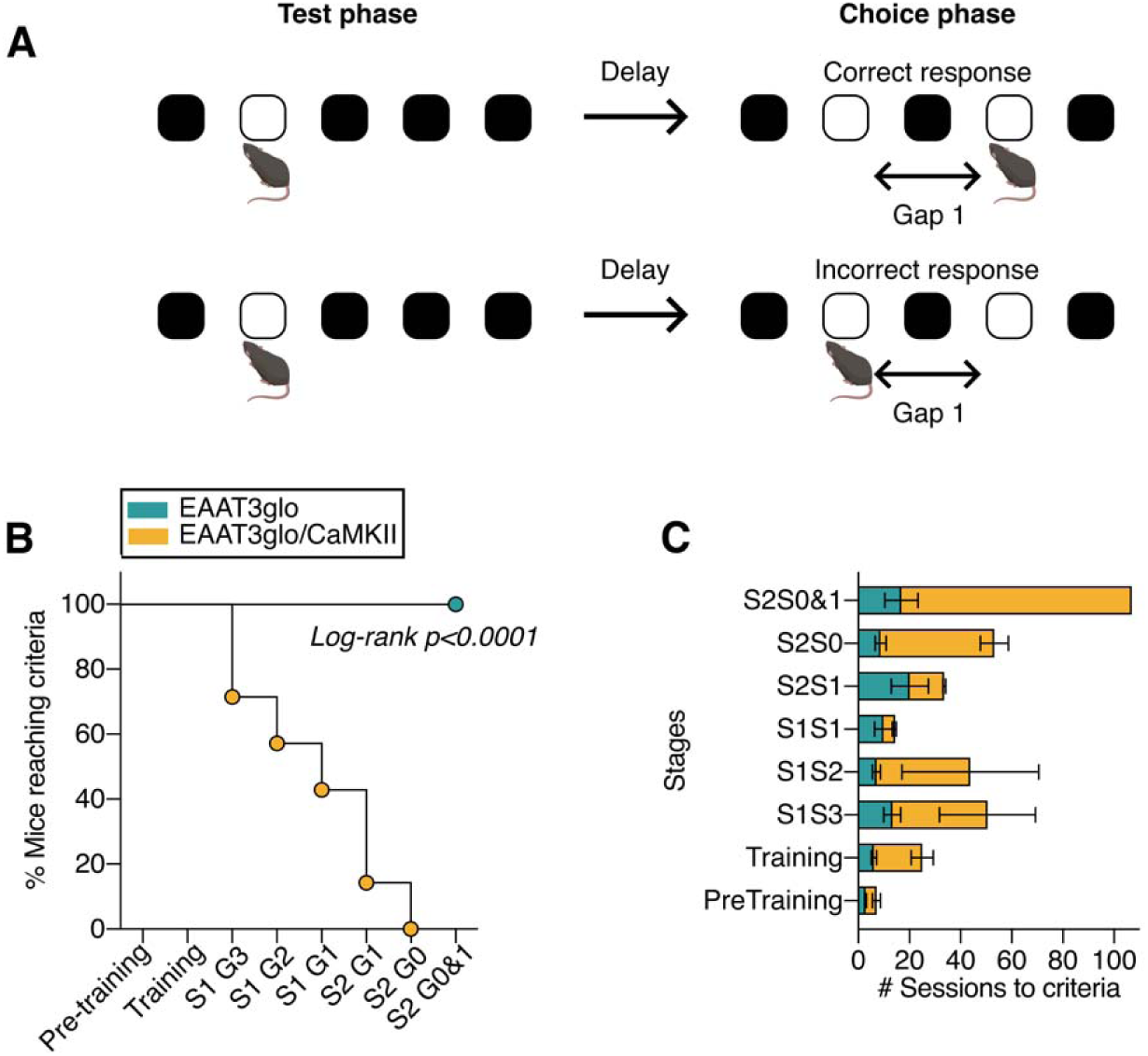
EAAT3glo/CaMKII mice fail to acquire the TUNL task. (A) Task schematic. (B) Percentage of mice reaching criterion and advancing to each TUNL stage. All EAAT3glo mice completed every stage (100%), while EAAT3glo/CaMKII mice dropped out progressively from training onwards, with none reaching S2S0 or S2S0&1. Log-rank (Mantel-Cox): χ²(1)=15.99, p<0.0001. Stage numbers: 1=Pre-training, 2=Training, 3=S1G3, 4=S1G2, 5=S1G1, 6=S2S1, 7=S2S0, 8=S2S0&1. (C) Sessions required to reach criterion at each stage, for animals that reached it. No significant between-genotype differences were found at any individual stage (Mann-Whitney, all p>0.05), due to the small n resulting from progressive dropout. n per stage for EAAT3glo/CaMKII: Pre-training n=7, Training n=6, S1G3 n=4, S1G2 n=3, S1G1–S2S0&1 n≤2. EAAT3glo n=7, EAAT3glo/CaMKII n=7. Mean ± SEM. ****p<0.0001.

To complement this finding with a simpler working memory measure, mice were tested in the T-maze spontaneous alternation task. Given that EAAT3glo/CaMKII mice failed to acquire the TUNL task — a paradigm that directly assesses working memory — we hypothesized *a priori* that these mice would show reduced spontaneous alternation compared to controls. Control mice (n=9, 2 outliers removed by sequential Grubbs test, α=0.05) showed a spontaneous alternation index significantly above chance level (one-sample t-test vs 50%: t(8)=2.857, p=0.021, Fig. 5A), while EAAT3glo/CaMKII mice (n=13) did not differ from chance (t(12)=0.210, p=0.837). Direct between-genotype comparison showed a trend toward reduced alternation in EAAT3glo/CaMKII mice (Mann-Whitney two-tailed: U=32, p=0.072). Based on the a priori hypothesis that EAAT3glo/CaMKII mice would show reduced alternation — derived from their failure to acquire the TUNL task — a one-tailed test was additionally performed (p=0.037). Choice latency did not differ between genotypes across trials (two-way ANOVA mixed model: F(6,120)=0.645, p=0.69, Fig. 5B), ruling out differences in decision speed or motor performance. Together, these results are consistent with a working memory impairment in EAAT3glo/CaMKII mice.

**Figure 5.**
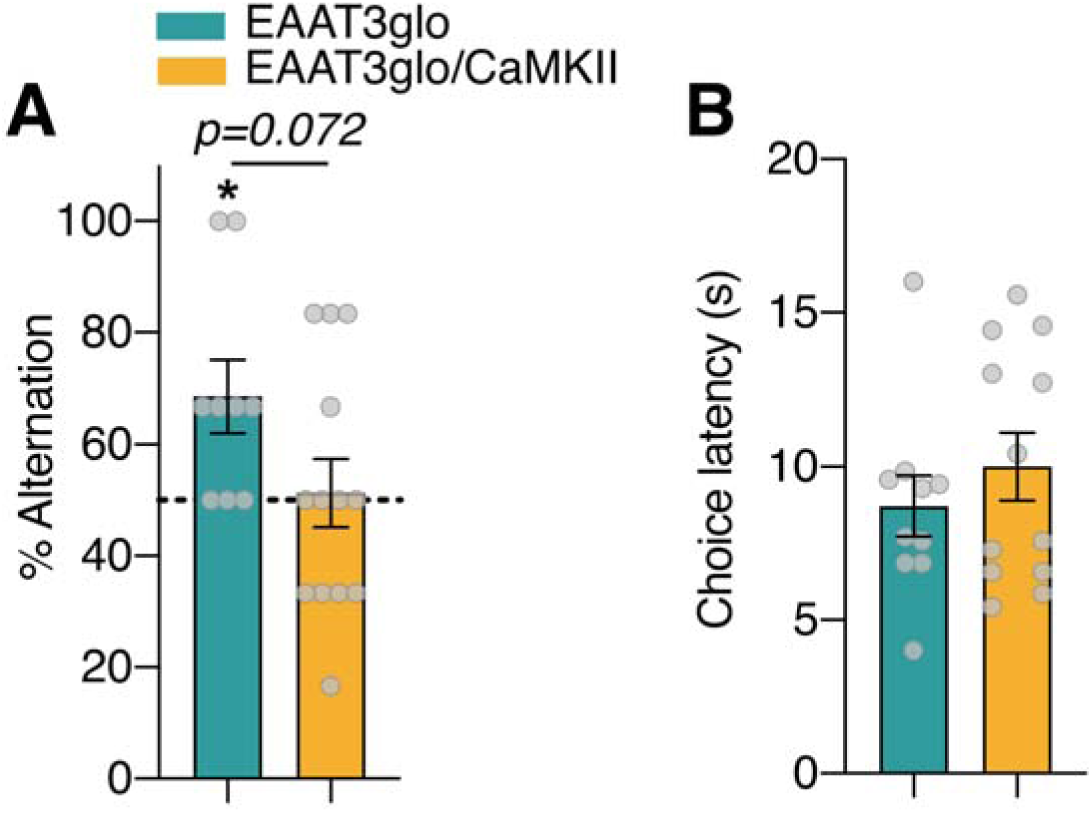
EAAT3glo/CaMKII mice show reduced spontaneous alternation in the T-maze. (A) EAAT3glo mice alternated above chance level (one-sample t-test vs 50%: t(8)=2.857, p=0.021), while EAAT3glo/CaMKII mice performed at chance (t(12)=0.210, p=0.837). Between-genotype comparison showed a trend toward reduced alternation in EAAT3glo/CaMKII mice (Mann-Whitney two-tailed: U=32, p=0.072; one-tailed based on a priori hypothesis: p=0.037). Dashed line indicates 50% chance level. (B) Choice latency did not differ between genotypes (F(6,120)=0.645, p=0.69), ruling out differences in speed or motor performance. EAAT3glo n=9 (2 outliers removed by Grubbs test, α=0.05), EAAT3glo/CaMKII n=13. Mean ± SEM with individual data points. *p<0.05.

## DISCUSSION

Here, we demonstrated the consequences of EAAT3 overexpression in the forebrain on executive functions. Using behavioral paradigms, we assessed performance across three domains commonly associated with executive function. While these domains are operationally useful for organizing our findings, we acknowledge that the tasks used in our study engage overlapping cognitive and neural processes, and that the behavioral alterations observed may reflect shared as well as distinct mechanisms. EAAT3glo/CaMKII mice display deficits in operant extinction and reversal, impaired working memory and impulsivity. Taken together, our data revealed a profile of alterations in executive function in a genetic mouse model that displays OCD-relevant phenotypes, highlighting a role for EAAT3 and the control of glutamate levels in the regulation of these executive functions.

EAAT3 is encoded by the *SLC1A1* gene, which was consistently associated in early OCD genetic studies (27, 28), although it has not yet reached the level of statistical significance required to be considered a definitive risk gene for OCD (29, 30). Despite the lack of association in more recent genome-wide association studies, some of the *SLC1A1* gene variants most associated with OCD reportedly increase *SLC1A1* expression in human postmortem brain as well as in cell models (31, 32), suggesting that increased EAAT3 expression/function may be linked to phenotypes that are relevant to OCD (33). This notion has been supported by several animal studies that implicate EAAT3 in abnormal repetitive behaviors, such as increased grooming and enhanced sensitivity to amphetamine-induced hyperlocomotion and stereotypy (16, 34–38).

EAAT3glo/CaMKII mice exhibit a profound deficit in operant extinction, extending prior observations of impaired long-term fear extinction (16, 39). This underscores a key deficit in cognitive flexibility in this model. Alterations in the glutamatergic system impact have been demonstrated to impact performance in various behavioral tests that assess executive functions in mice. For instance, the administration of NMDA receptor antagonists has been shown to impair performance on a variety of cognitive tasks, including TUNL (18, 40, 41), cognitive flexibility (42–44) and inhibitory control (5-CSRTT) (45). Interestingly, EAAT3glo/CaMKII mice have altered subunit composition and function of NMDA receptors at cortico-striatal synapses, (16), suggesting that alterations in NMDA receptors might underline the alterations observed in executive function. Of note, studies using another mouse model relevant for OCD, which is deficient in the glutamatergic synapse scaffolding protein SAPAP3 (SAPAP3 KO mice), reported that they can acquire Pavlovian learning but exhibit cognitive inflexibility by failing to adjust their conditioned approach behavior (46–48). However, SAPAP3 KO mice do not show deficits in the extinction of Pavlovian or operant learning (49). Together, these studies support a role for the glutamatergic signaling in the pathophysiology of OCD, particularly in different manifestations of executive dysfunction (20, 50).

The impaired extinction seen in EAAT3glo/CaMKII mice model strongly suggests the presence of executive dysfunction. Extinction learning, whether operant or Pavlovian, involves the formation of a new, competing learning process (51–54). The formation of this new memory requires setting aside an already acquired rule for a new one and engaging in goal-directed behavior, a process that critically depends on executive functions (55). The glutamatergic system plays a key role in memory extinction. For instance, the administration of the glutamatergic agonist D-Cycloserine has been reported to facilitate extinction of operant learning in mice (56, 57). Similarly, the administration of an mGluR5 antagonist attenuates instrumental extinction in a mouse model of cocaine self-administration, suggesting that mGluR5 plays an essential role in extinction learning (58). Furthermore, the core neural circuit associated with the extinction process comprises the mPFC, amygdala, hippocampus and thalamus, which are predominantly interconnected via glutamatergic pathways (53, 59).

Glutamatergic NMDAR agonists reportedly facilitate fear extinction (60, 61) and NMDAR activation in the infralimbic cortex appears to mediate the increase in glutamate neurotransmission during the consolidation of extinction learning (60). Likewise, it has been proposed that both fear memory extinction and its consolidation, which require precise control of glutamate levels, are altered in OCD, resulting in increased compulsive behaviors as an attempt to “get rid of” the fear (and its triggers) experienced by individuals with OCD (19). In this sense, our results contribute to the OCD study field by highlighting the behavioral changes that occur when manipulating the expression and activity levels of EAAT3, one of the regulators of glutamate levels in the brain. Accordingly, clinical analyses have demonstrated that a decrease in glutamate levels in the ventromedial prefrontal cortex (vmPFC) correlates with poorer performance in fear memory extinction tests in OCD individuals (62). Further supporting the role of glutamate levels, other studies have shown that N-Acetylcysteine, an effective glutamate-modulating drug, can be utilized as a therapeutic option for patients with moderate-to-severe OCD (63). Similarly, the effectiveness of transcranial magnetic stimulation therapy directly correlates with increases in glutamate levels in the striatum of patients (64). Of note, a recent meta-analysis demonstrated that various glutamatergic treatments, including riluzole, amantadine, D-cycloserine, dextromethorphan, lamotrigine, L-carnosine, memantine, and N-acetylcysteine, can significantly reduce OCD symptoms in patients (65). Although literature is mixed with reported increase or decrease of glutamate levels in different brain areas of OCD patients, our data prompts us to hypothesize that EAAT3 overexpression may be decreasing local glutamate levels, thereby impairing memory extinction mechanisms. Further studies to determine the differential expression of EAAT3 in different brain areas, particularly in subareas of the cortex (i.e., PFC vs AAC), and its implication in glutamate levels and cognitive processes are required in animal models relevant for OCD, as we propose in the EAAT3glo/CaMKII mouse model.

We also show that EAAT3glo/CaMKII mice exhibit impaired operant reversal learning, failing to adapt to the new rule. This reflects a cognitive flexibility impairment in operant learning, which parallels the above-mentioned clinical data. Recently, Petroccione et al. (66) described that EAAT3 modulates the excitability of M1-MSN neurons in the DLS, which has implications for behavioral flexibility in mice. Although these studies reinforce the idea that glutamatergic system alterations are associated with executive dysfunction in compulsivity models, we cannot attribute deficits in these reverse learning tasks solely to an alteration in cognitive flexibility. While these results likely stem from the inability of EAAT3glo/CaMKII mice to abandon the previously learned rule, perseverative behaviors or an inability to remember the new rule may also contribute to these outcomes. To address this, we employed the 5-CSRTT and TUNL tests, which allow the isolation of different executive function components. The 5-CSRTT assesses inhibitory control, while the TUNL test assesses working memory. These tests were conducted in Bussey-Saksida touchscreen operant chambers, which facilitated a more detailed analysis of animal behavior and a more accurate isolation of one executive function from another, adding high translational relevance to human tests (24).

In the 5-CSRTT, premature responses (i.e., made before the stimulus appears) are the main measure of waiting impulsivity under standard testing conditions and are regulated by prefrontal-accumbal circuits (23, 67, 68). EAAT3^glo^/CaMKII mice showed more premature responses than controls at baseline, indicating increased waiting impulsivity. Since EAAT3glo/CaMKII mice have reportedly altered glutamatergic transmission at corticostriatal synapses (16), it is plausible that these synaptic impairments contribute to the increased impulsivity observed in this model. To more directly test inhibitory control, additional challenge sessions were run varying the inter-trial intervals (ITI), in order to remove the animal’s ability to anticipate when the stimulus will appear, making response suppression genuinely necessary (24, 69, 70). Under these conditions, no differences were found between genotypes, suggesting that the baseline increase in premature responses may not directly reflect a true deficit in inhibitory control, but rather a difference in how animals estimate the timing of the interval; further studies using temporal discrimination tasks would be needed to confirm this.

Working memory deficits have also been reported in OCD (12). Our results suggest that mice overexpressing EAAT3 exhibit significantly poorer performance in the acquisition phase of the TUNL task, which is used to evaluate working memory and pattern separation. EAAT3glo/CaMKII mice do not meet the criteria to complete the acquisition phase, which precludes them from advancing to the testing phase. TUNL acquisition critically depends on hippocampal and prefrontal circuits and is sensitive to NMDA receptor hypofunction (18, 40, 71). Moreover, EAAT3glo/CaMKII mice failed to meet the alternation criterion in the T-maze. Our findings agree with a previous report of altered spontaneous alternation in another animal model of compulsivity (72), suggesting a deficit in working memory and/or cognitive flexibility. The equivalent choice latency observed between genotypes in the T-maze rules out differences in motor performance or decision speed as alternative explanations. Further experiments should directly evaluate if the alterations in NMDA receptor subunit composition and impaired corticostriatal synaptic plasticity reported in EAAT3glo/CaMKII mice (16) also occur in prefrontal and hippocampal regions, which would allow to unravel the mechanisms underlying the impaired working memory in this model.

In summary, our findings demonstrate that increased expression of EAAT3 is accompanied by executive dysfunction, impacting cognitive flexibility, working memory and impulsivity, while preserving attention and inhibitory control. These deficits observed in EAAT3glo/CaMKII mice provides a useful basis for dissecting the contribution of glutamatergic mechanisms to specific executive functions and phenotypes that are relevant to OCD and other disorders, offering insights into the mechanistic underpinnings as well as to the heterogeneity of this disorder.

## DATA AVAILABILITY

Data are available upon request to the corresponding authors.

## AUTHOR CONTRIBUTIONS

FHB contributed to acquisition, analysis and interpretation of data, drafting and editing the manuscript. MCM contributed to acquisition, analysis and interpretation of data and editing the manuscript. WPB contributed to acquisition and analysis of data and editing the manuscript. PAH contributed to acquisition and analysis of data and editing the manuscript. MJC contributed to acquisition of data and editing the manuscript. AEC contributed to the experimental conceptualization and experimental design, and editing the manuscript. JPC contributed to the experimental conceptualization and experimental design, analysis and interpretation of data, drafting and editing the manuscript. PRM contributed to the experimental conceptualization and experimental design, analysis and interpretation of data, drafting and editing the manuscript. All authors approved the submitted version of this manuscript.

(A) Time course of acquisition of operant behavior in EAAT3glo and EAAT3glo/CMKII mice. Similar increase in active responses (B), decrease in inactive (C) and unaffected center responses (D) were observed in both genotypes (N= 22 EAAT3glo, 25 EAAT3glo/CMKII) between days 1 vs 9 of the protocol. (E) Time course of extinction. EAAT3glo mice showed a decrease in active (F), inactive (G) and center responses (H), while EATT3glo/CaMKII mice did not show differences between days 1 vs 6 of extinction protocol (N= 11 EAAT3glo, 11 EAAT3glo/CMKII). Data represent Mean+/-s.e.m. *P < 0.05; ***P < 0.001; ****P< 0.0001

