## Supplementary Material for "Executive Dysfunctions In The EAAT3 Overexpressing Mouse Model Of Compulsive Behavior"

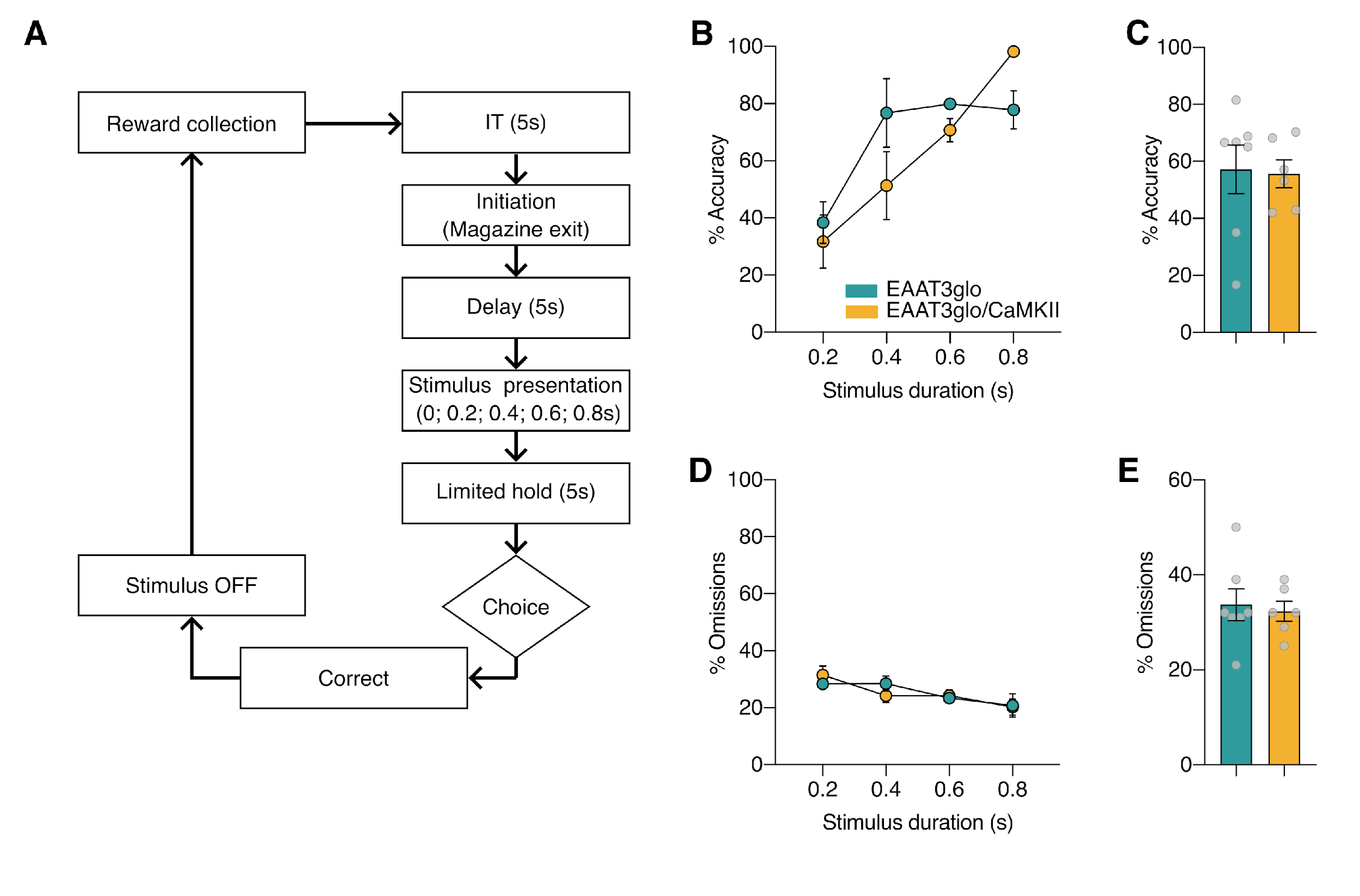

**Supplementary figure 1. EAAT3glo/CMKII mice do not show alterations in 5-CSRTT – Short ITI performance.**

EAAT3glo/CMKII mice and control EAAT3glo littermates underwent evaluation in the 5-CSRTT. (A) 5-CSRTT -short ITI flux diagram, (B) reward collection latency, (C) accuracy, (D) omission percentage, (E) perseverative responses percentage, and (F) premature responses percentage in both EAAT3glo and EAAT3glo/CMKII mice. (N= EAAT3glo: 8; EAAT3glo/CamKII: 6; (Mean+/-s.e.m))

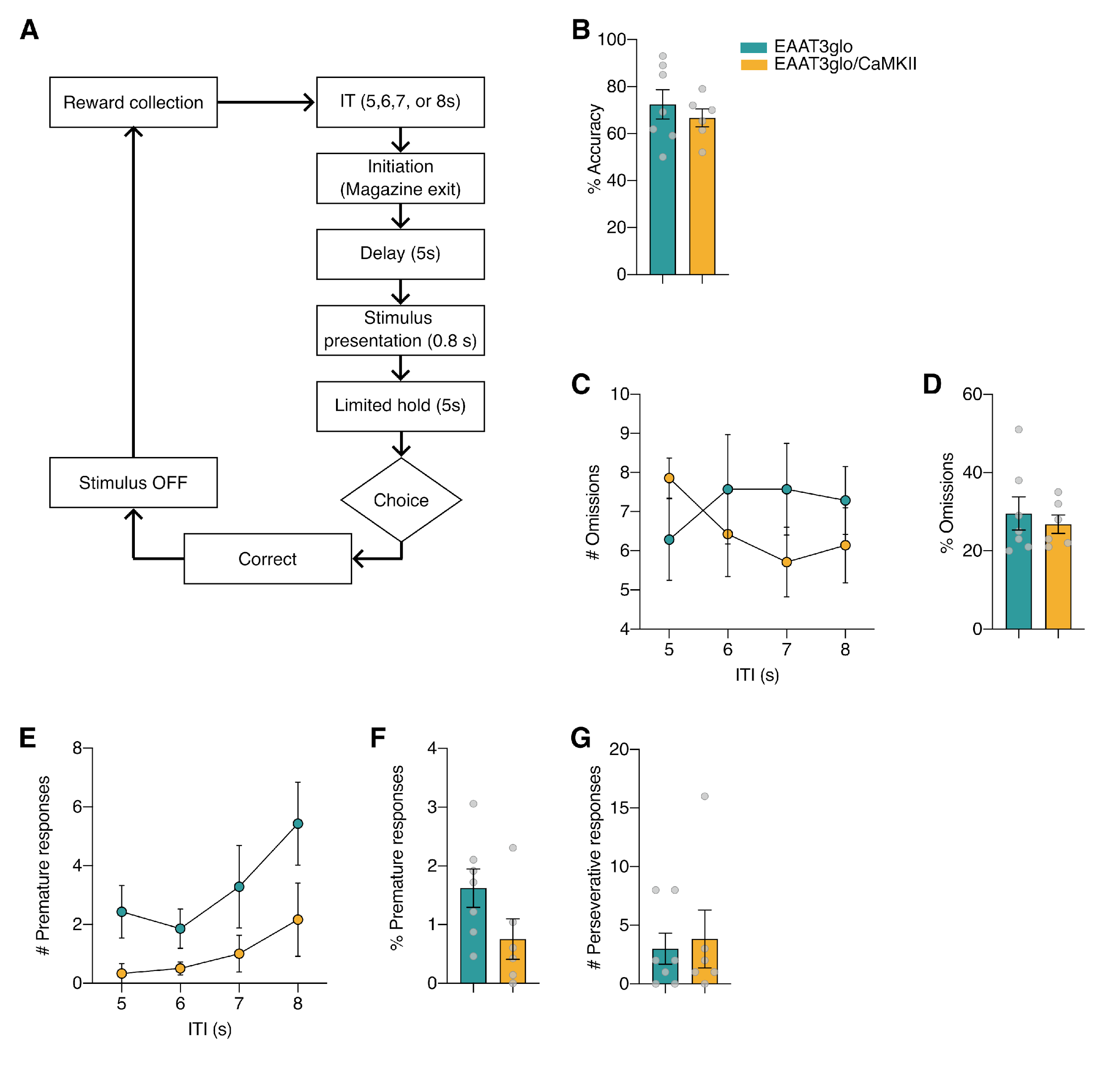

**Supplementary figure 2. EAAT3glo/CMKII mice do not show alterations in 5-CSRTT – long ITI performance.**

EAAT3glo/CMKII mice and control EAAT3glo littermates underwent evaluation in the 5CSRTT using Bussey & Saksida operant chambers. (A) 5-CSRTT -long ITI flux diagram, (B) reward collection latency, (C) accuracy, (D) omission percentage, (E) perseverative responses percentage, and (F) premature responses percentage in both EAAT3glo and EAAT3glo/CMKII mice. (N= EAAT3glo: 8; EAAT3glo/CamKII: 6; (Mean+/-s.e.m))

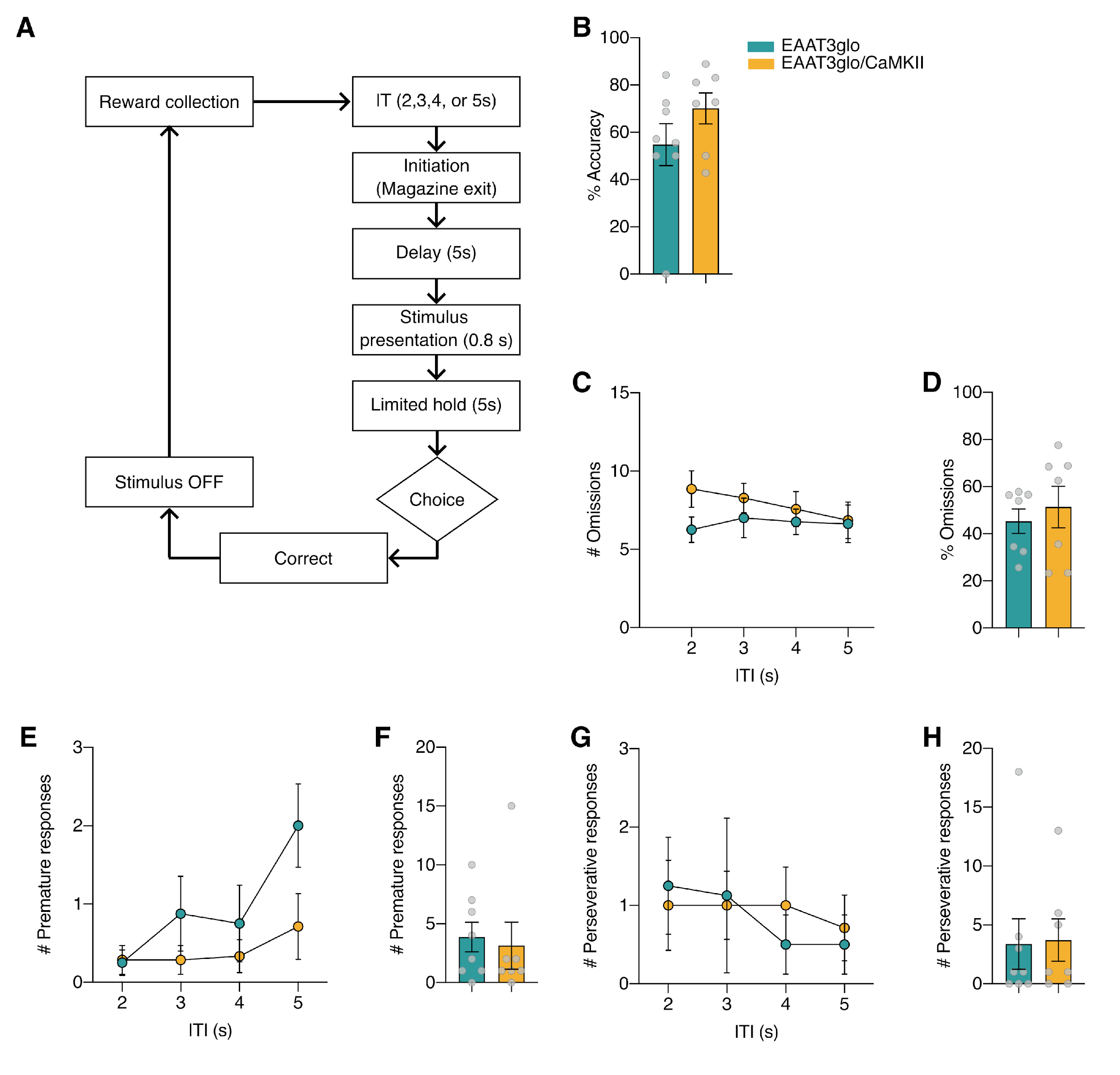

**Supplementary figure 3. EAAT3glo/CMKII mice do not show alterations in 5-CSRTT – variable stimulus ITI performance.**

EAAT3glo/CMKII mice and their EAAT3glo littermates underwent evaluation in the 5CSRTT using Bussey & Saksida operant chambers. (A) 5-CSRTT -variable stimulus flux diagram, (B) reward collection latency, (C) accuracy, (D) omission percentage, (E) perseverative responses percentage, and (F) premature responses percentage in both EAAT3glo and EAAT3glo/CMKII mice. (N= EAAT3glo: 8; EAAT3glo/CamKII: 6; (Mean+/-s.e.m))

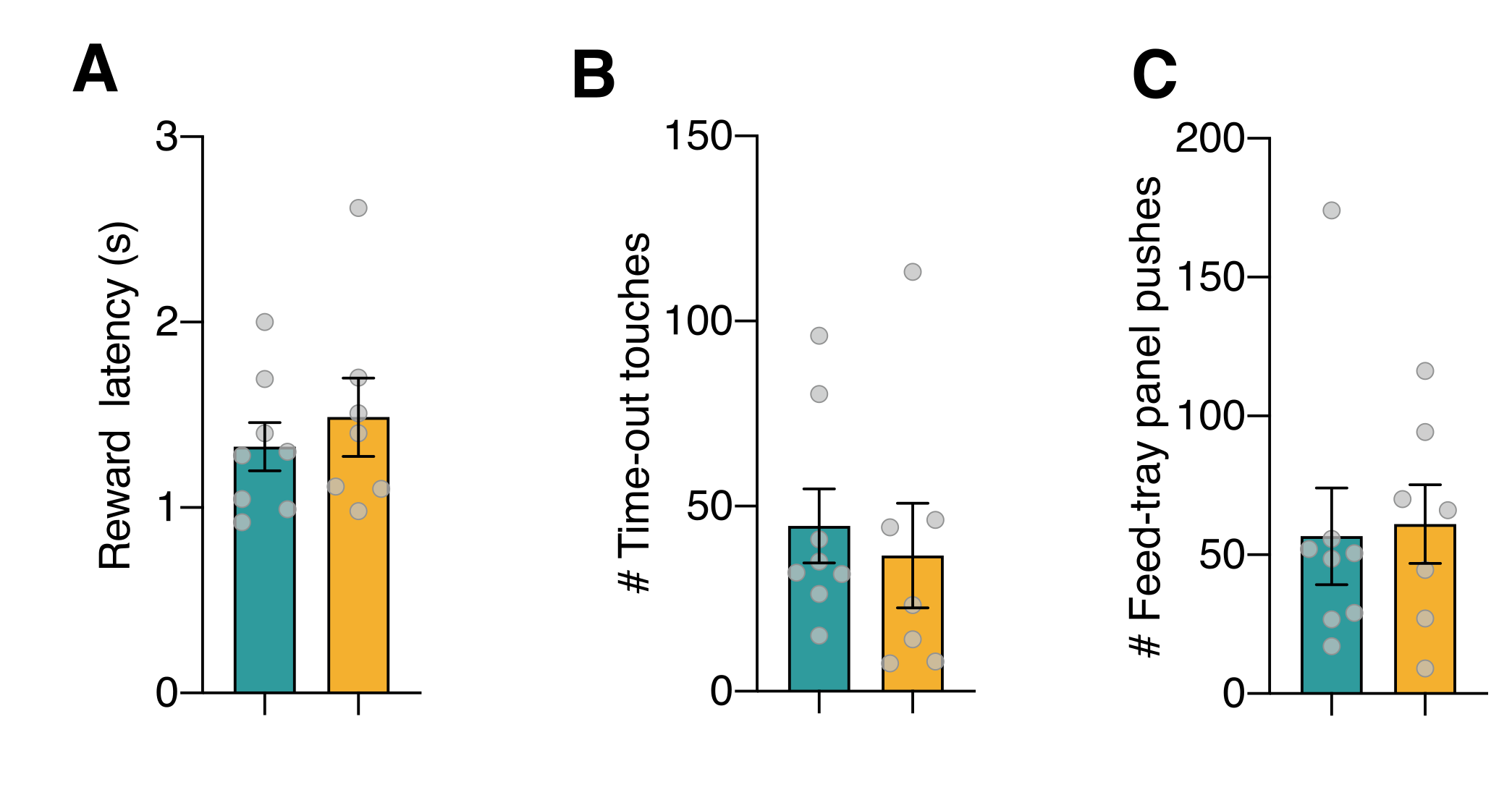

**Supplementary figure 4. Basic 5-CSRTT performance controls**. No significant between-genotype differences were observed in reward collection latency (A; Mann-Whitney, p=0.55), time-out touches (B; Mann-Whitney, p=0.48), or feed-tray panel pushes (C; Mann-Whitney, p=0.61), confirming equivalent motivation and general motor activity across groups. EAAT3glo n=8, EAAT3glo/CaMKII n=6. Mean ± SEM with individual data points.

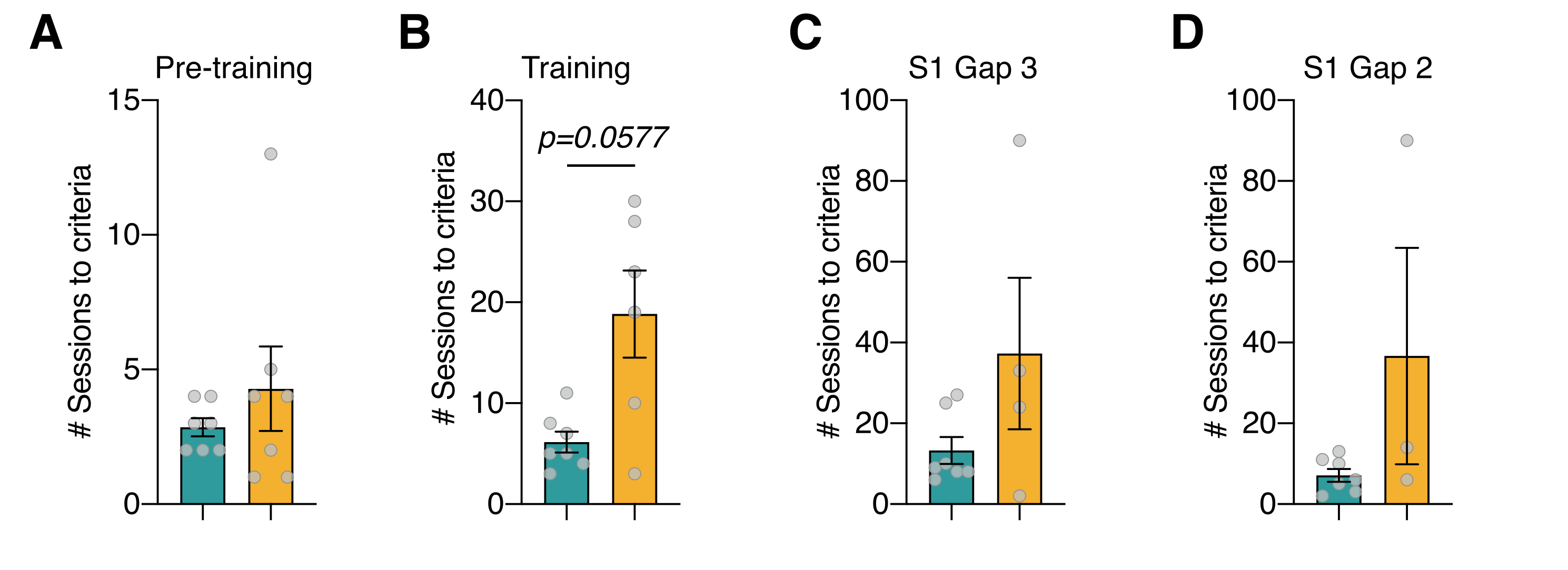

**Supplementary figure 5. Sessions required to reach criterion at each TUNL stage, for animals that completed that stage.** (A) Pre-training: no significant differences between genotypes (Mann-Whitney, p>0.05). (B) Training: EAAT3glo/CaMKII mice tended to require more sessions than controls (Mann-Whitney, p=0.058). (C) S1 Gap 3: no significant differences (Mann-Whitney, p=0.391). (D) S1 Gap 2: no significant differences (Mann-Whitney, p=0.137). Data shown as mean ± SEM with individual data points. Note that n per stage decreases progressively for EAAT3glo/CaMKII due to dropout: Pre-training n=7, Training n=6, S1 Gap 3 n=4, S1 Gap 2 n=3. EAAT3glo n=7 for all stages.

**Supplementary Table 1**

| Phase | Description | Duration | Criteria |
| --- | --- | --- | --- |
| Habituation 1 | The mouse is left in the chamber for a 10-minute session, with all lights off. No stimuli or rewards are presented. The mouse's activity is monitored. | 1 day | -------- |
| Habituation 2a | The mouse is left in the chamber for a 20-minute session with the light on. 150 uL of reward is added to the food tray. Once the mouse enters and leaves the food tray, the tray light is turned off. There is a 10-s delay before the tray light is turned on and reward is delivered again. The procedure is repeated until the session ends. | 1 day | -------- |
| Habituation 2b | The mouse is left in the chamber for a 40-minute session with the light on. 150 uL of reward is added to the food tray. Once the mouse enters and leaves the food tray, the tray light is turned off. There is a 10-s delay before the tray light is turned on and reward is delivered again. The procedure is repeated until the session ends. | 1 day | -------- |
| Habituation 2c | The mouse is left in the chamber for a 60-minute session with the light on. 150 uL of reward is added to the food tray. Once the mouse enters and leaves the food tray, the tray light is turned off. There is a 10-s delay before the tray light is turned on and reward is delivered again. The procedure is repeated until the session ends. | 1-2 days | The animal must consume the entire reward |
| Initial Touch Training (Pre-training in TUNL) | The stimulus (a white square) is randomly displayed in one of the five windows. After a delay, the image is removed and the reward is delivered. Food delivery is accompanied by the illumination of the tray light and an optional tone. When the animal goes to collect the reward, the stimulus is displayed in another location. If the mouse touches the screen while the stimulus is displayed, it is removed, a tone is played, and the reward is delivered. Collecting the reward triggers the start of a new trial. | 1-3 days | Complete 30 essays in 30 minutes |
| Must Touch Training (Training in TUNL) | The stimulus (a white square) is randomly displayed in one of the 5 windows. The mouse must touch the stimulus to obtain a tone/food response. There is no response if the mouse touches a blank portion of the screen. Reward delivery is accompanied by the illumination of the tray light and a tone. The entry to collect the reward turns off the tray light and the ITI begins. After the ITI period (default 5 s), another image is displayed. | 1-7 days | Complete 20 essays in 30 minutes |

**Supplementary Table 2**

| **Separation level** | **Stage 1**  (No stimuli in center) | **Stage2**  (Halfway through the assay with a sample stimulus in center) |
| --- | --- | --- |
| S3 | 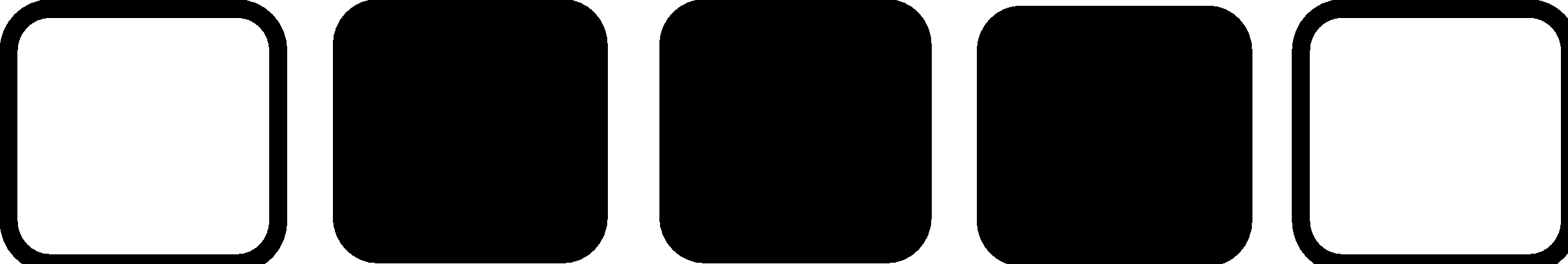 |  |
| S2 | 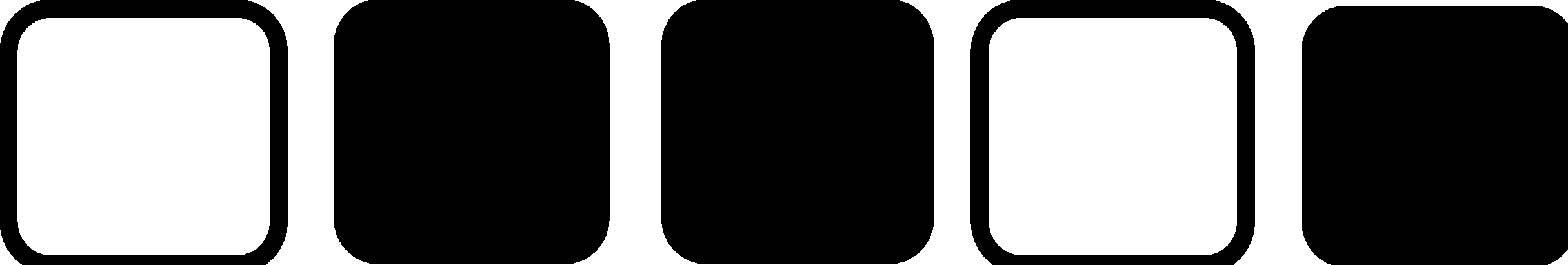  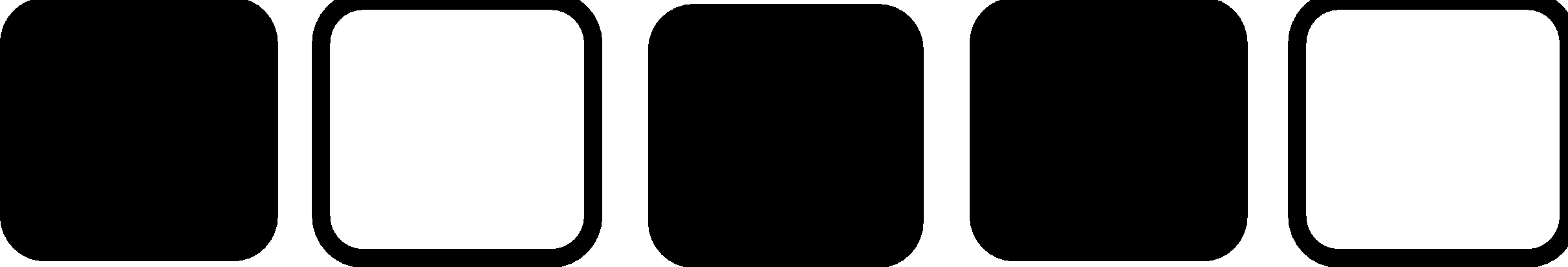 |  |
| S1 | 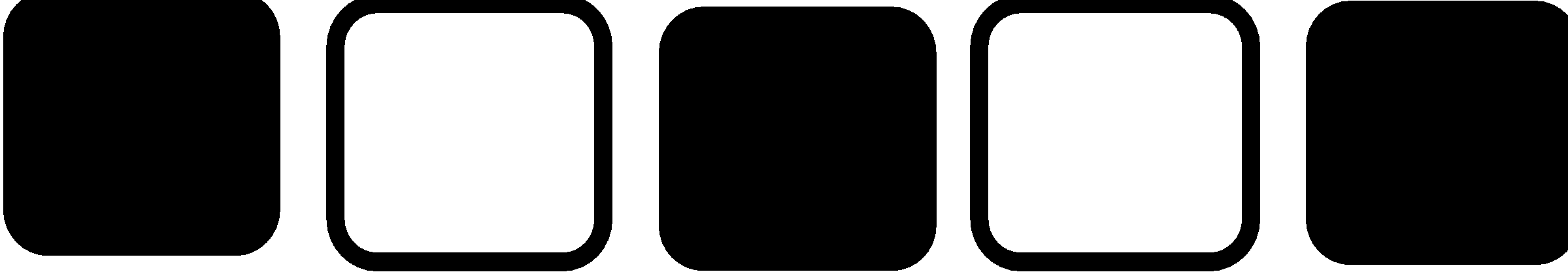 | 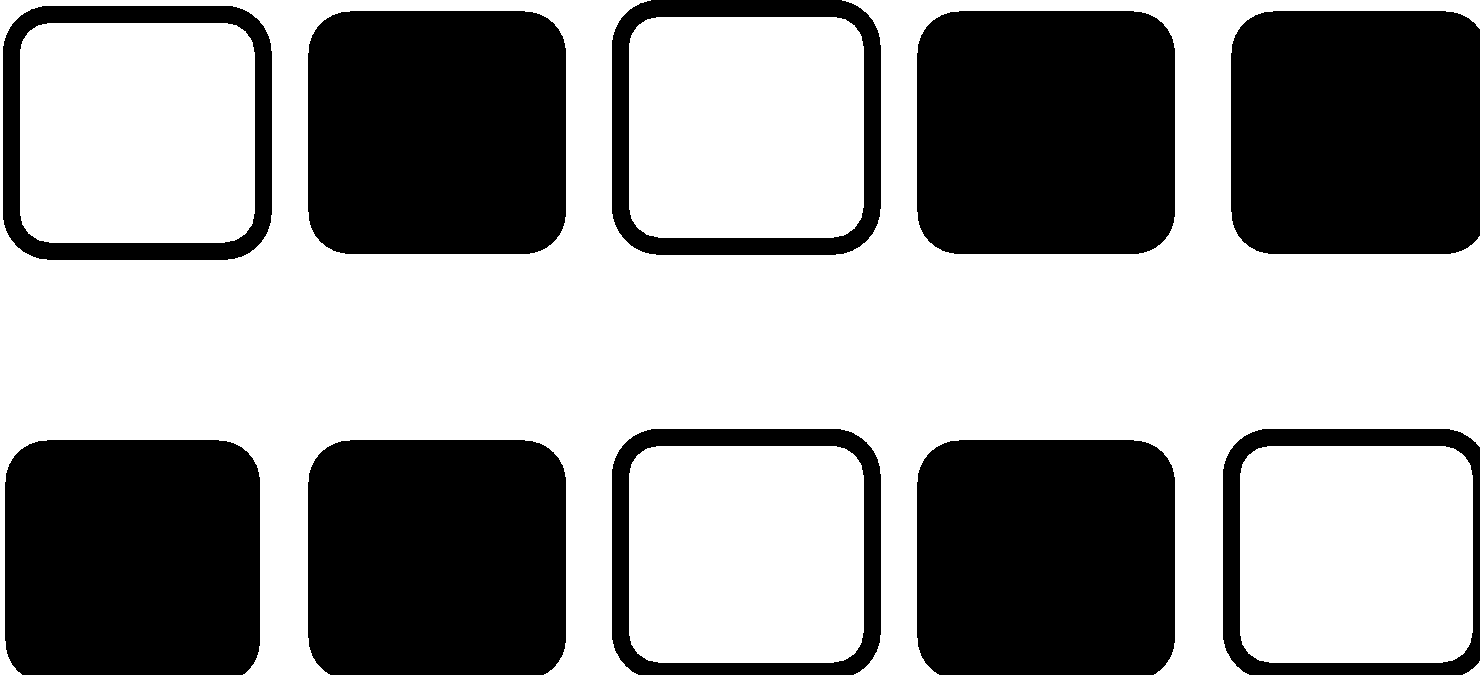 |
| S0 |  | 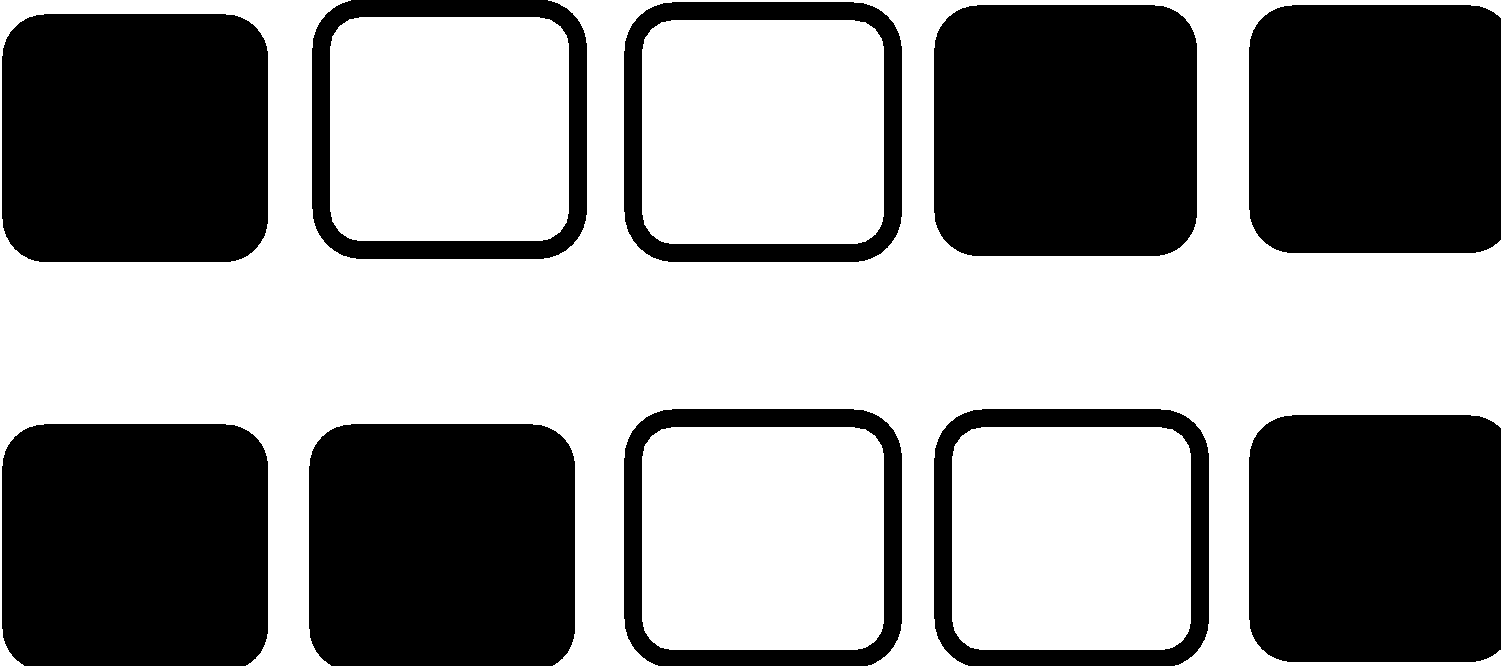 |

**Supplementary Table 3. Summary of statistical analyses**

All analyses: GraphPad Prism 8.4.3. α = 0.05. ns = not significant; * p<0.05; *** p<0.001; **** p<0.0001; trend = p<0.10.

**1. Acquisition of operant behavior**

*Mixed-effects REML (Matching: Stacked). EAAT3glo n=22, EAAT3glo/CaMKII n=25.*

| **Measure** | **Effect** | **F statistic** | **df** | **p value** | **Significance** |
| --- | --- | --- | --- | --- | --- |
| Active hole | Session × Genotype | F=0.671 | 8, 182 | p=0.715 | ns |
| Active hole | Session | F=8.203 | 8, 182 | p<0.0001 | **** |
| Inactive hole | Session × Genotype | F=0.654 | 8, 397 | p=0.732 | ns |
| Inactive hole | Session | F=12.45 | 8, 397 | p<0.0001 | **** |
| Magazine | Session × Genotype | F=0.791 | 8, 200 | p=0.611 | ns |

*Day 1 vs Day 9 within-genotype comparisons.*

| **Measure** | **Genotype** | **p value (Day 1 vs 9)** | **Significance** |
| --- | --- | --- | --- |
| Active hole | EAAT3glo | p<0.0001 | **** |
| Active hole | EAAT3glo/CaMKII | p<0.001 | *** |
| Inactive hole | EAAT3glo | p<0.05 | * |
| Inactive hole | EAAT3glo/CaMKII | p<0.001 | *** |
| Magazine | EAAT3glo | p=0.15 | ns |
| Magazine | EAAT3glo/CaMKII | p=0.15 | ns |

**2. Extinction of operant behavior**

*Mixed-effects REML (Matching: Stacked). EAAT3glo n=11, EAAT3glo/CaMKII n=11.*

| **Measure** | **Effect** | **F statistic** | **df** | **p value** | **Significance** |
| --- | --- | --- | --- | --- | --- |
| Active hole | Session × Genotype | F=2.884 | 5, 100 | p=0.018 | * |
| Active hole | Session | F=12.50 | 5, 100 | p<0.0001 | **** |
| Inactive hole | Session × Genotype | F=2.106 | 5, 50 | p=0.080 | trend |
| Magazine | Session × Genotype | F=0.405 | 5, 105 | p=0.831 | ns |

*AUC between-genotype comparisons (Mann-Whitney U test).*

| **Measure** | **p value (AUC)** | **Significance** |
| --- | --- | --- |
| Active hole AUC | p=0.176 | ns |
| Inactive hole AUC | p=0.046 | * |
| Magazine AUC | p=0.451 | ns |

**3. Reversal learning**

*Mixed-effects REML (Matching: Stacked). Post-hoc Sidak between genotypes at individual sessions: all ns. EAAT3glo n=11, EAAT3glo/CaMKII n=11.*

| **Measure** | **Effect** | **F statistic** | **df** | **p value** | **Significance** |
| --- | --- | --- | --- | --- | --- |
| -CS hole | Session × Genotype | F=3.398 | 3, 83 | p=0.023 | * |
| +CS hole | Session × Genotype | F=3.509 | 3, 83 | p=0.020 | * |
| Magazine | Session × Genotype | F=0.226 | 3, 66 | p=0.878 | ns |

*AUC between-genotype comparisons (Mann-Whitney U test).*

| **Measure** | **p value (AUC)** | **Significance** |
| --- | --- | --- |
| -CS hole AUC | p=0.182 | ns |
| +CS hole AUC | p=0.038 | * |
| Magazine AUC | p=0.451 | ns |

**4. 5-Choice Serial Reaction Time Task (5-CSRTT)**

*Variance equality assessed by F-test prior to comparisons. EAAT3glo n=8, EAAT3glo/CaMKII n=6.*

| **Measure** | **Test** | **Statistic** | **p value** | **Significance** |
| --- | --- | --- | --- | --- |
| Premature responses | Welch's t-test (F-test p=0.010) | t(6.94)=2.670 | p=0.032 | * |
| Perseverative responses | Mann-Whitney | — | p=0.92 | ns |
| Response accuracy | Mann-Whitney | — | p=0.77 | ns |
| Omission rate | Mann-Whitney | — | p=0.93 | ns |
| Reward latency | Mann-Whitney | — | p=0.55 | ns |
| Time-out touches | Mann-Whitney | — | p=0.48 | ns |
| Feed-tray panel pushes | Mann-Whitney | — | p=0.61 | ns |

*One-day challenge tests (short ITI, long ITI, variable stimulus): no significant between-genotype differences (Figs. S1-S3).*

**5. T-maze spontaneous alternation**

*2 outliers removed from EAAT3glo by sequential Grubbs test (α=0.05). EAAT3glo n=9, EAAT3glo/CaMKII n=13.*

| **Measure** | **Test** | **Statistic** | **p value** | **Significance** |
| --- | --- | --- | --- | --- |
| Alternation index — EAAT3glo vs 50% | One-sample t-test | t(8)=2.857 | p=0.021 | * |
| Alternation index — EAAT3glo/CaMKII vs 50% | One-sample t-test | t(12)=0.210 | p=0.837 | ns |
| Between-genotype (two-tailed) | Mann-Whitney | U=32 | p=0.072 | trend |
| Between-genotype (one-tailed, a priori) | Mann-Whitney | U=32 | p=0.037 | * |
| Choice latency | Two-way ANOVA mixed model | F(6,120)=0.645 | p=0.69 | ns |

*One-tailed test based on a priori hypothesis derived from TUNL results (see Statistical Analysis in Methods).*

**6. Touchscreen TUNL task**

*EAAT3glo n=7, EAAT3glo/CaMKII n=7. n per stage for KO: Pre-training=7, Training=6, S1G3=4, S1G2=3, S1G1-S2S0&1 ≤2.*

| **Measure** | **Test** | **Statistic** | **p value** | **Significance** |
| --- | --- | --- | --- | --- |
| Survival curve (% reaching criteria) | Log-rank (Mantel-Cox) | χ²(1)=15.99 | p<0.0001 | **** |
| Sessions to criteria — Pre-training | Mann-Whitney | — | p>0.05 | ns |
| Sessions to criteria — Training | Mann-Whitney | — | p=0.058 | trend |
| Sessions to criteria — S1 Gap 3 | Mann-Whitney | — | p=0.391 | ns |
| Sessions to criteria — S1 Gap 2 | Mann-Whitney | — | p=0.137 | ns |
| Sessions to criteria — S1G1 onwards | — | — | n insufficient | — |

*Between-genotype comparisons at individual stages have limited power due to progressive dropout. Log-rank test on the full survival curve is the primary analysis.*
